# Dengue infection elicits skin tissue-resident and circulating CD8^+^ T-cells associated with protection from hospitalization

**DOI:** 10.1101/2025.07.11.664360

**Authors:** Noor Zayanah Hamis, Justin SG Ooi, Ka-Wai Cheung, Valerie Chew, Michaela Gregorova, Eugenia Ong Ziying, Kuan Rong Chan, Tun-Linn Thein, Yee-Sin Leo, David Chien Boon Lye, Eng Eong Ooi, Laura Rivino

## Abstract

Dengue is spreading globally and there is urgent need to define immune correlates of protection against this disease. Immune responses against dengue viruses have been studied in blood samples of dengue patients. However, dengue virus infection first occurs in the skin following the bite of an infected *Aedes* mosquito, and immune responses are initiated within this site. In this study, we investigated the phenotypic, functional and transcriptional profiles of skin and blood T-cell responses and their role in immunity in 73 dengue patients and 10 healthy volunteers. We show that the skin T-cell compartment undergoes dramatic reshaping compared to the blood of dengue patients. CD4^+^ and CD8^+^ T-cell responses were highly enriched in the skin compared to the blood of the same patients, and skin-based T-cells expressed markers associated with tissue-resident T (T_RM_) cells. While the magnitude of the CD4^+^ T-cell response in the skin was independent to that in the blood, CD8^+^ T-cell responses in skin and blood were positively correlated. Activated CD8^+^ T-cells in the skin expressed a core transcriptional signature of T_RM_ cells, further supporting their differentiation to the T_RM_ lineage during infection. The magnitude of both skin and blood CD8^+^ T-cell responses was associated with protection from hospitalization in this cohort. These data collectively support a protective role of skin-resident and circulating CD8^+^ T-cells in dengue and provide insights into the biology of T_RM_ cells in human infection. Our findings warrant evaluation of vaccination strategies that induce T_RM_ cells in the skin to enhance protection against dengue.

**One Sentence Summary:** Dengue infection elicits skin tissue-resident and circulating CD8^+^ T-cells associated with protection from hospitalization in adult dengue patients.

## Introduction

Dengue is caused by dengue virus (DENV), a mosquito-borne flavivirus estimated to infect 390 million people, causing 300,000 severe dengue cases and 20,000 deaths per year(*1*). Urbanization and human mobility, both of which are now further complicated by climate change, have driven a rapid, approximately 10-fold increase in dengue cases in the last 20 years(*2*). DENV co-circulates as four genetically distinct infectious serotypes (DENV 1-4); infection with any DENV serotype may be asymptomatic or cause a range of clinical manifestations from a self-limiting febrile illness to severe dengue characterised by hypovolemic shock from uncontrolled plasma leakage, internal haemorrhage, and organ dysfunction(*3*). The pathogenesis of dengue is multifactorial and poorly understood, although altered host immune responses are believed to play key roles(*3, 4*). No licensed therapy exists for dengue. There are two licensed dengue vaccines which show unbalanced protection towards DENV1-4 despite their ability to elicit neutralizing antibodies against all four DENV serotypes, thus highlighting our incomplete understanding of the immune correlates of protection for dengue(*5*). In particular, the contribution of T-cells for protective immunity to DENV remains unclear. Studies to date have focused on analyses of T-cells in the blood of dengue patients which may not be representative of T-cell responses occurring within tissues where the majority of T-cell responses occur during infection.

T-cell responses initiate in the skin following the bite of a DENV-infected *Aedes* mosquito. We and others have shown that T-cell priming in dengue occurs in the skin/skin-draining lymph-nodes(*6*), where DENV-specific T-cells acquire expression of the skin-homing receptor cutaneous leukocyte-associated antigen (CLA)(*7*). CLA binds to E-selectin (CD62E) expressed on dermal endothelial cells and facilitates T-cell homing to the skin(*8*). It remains unknown whether DENV-specific T-cells in the skin differentiate into tissue-resident T-cells (T_RM_) and whether these play a role in protective immunity to dengue.

T_RM_ cells were identified a decade ago as specialised T-cell subsets located within tissues, which display superior protective efficacy towards pathogens compared to their circulating counterparts(*9, 10*). T_RM_ cells clear virus-infected cells through release of cytotoxic granules and cytokines/chemokines, the latter also acting as signals to recruit other leukocytes to the tissue(*11, 12*). In mice, T_RM_ cells were shown to have a key protective role against herpes simplex virus (HSV) in the skin, and influenza and respiratory syncytial viruses (RSV) in the lungs, with dispensable roles of circulating T-cells (*13, 14*). The role of T_RM_ cells in human infection remains less clear due to the difficulties in accessing the human tissues where these cells reside. T_RM_ cells were detected in human acute and chronic viral infections (Influenza, RSV, Hepatitis B and Human Immunodeficiency viruses)(*14–16*), with some studies reporting an association of T_RM_ cell frequencies with enhanced viral control(*17*). Understanding the contribution of T_RM_ cells to protective immunity is critical for evaluation of vaccine delivery routes that could improve vaccine efficacy by boosting generation of protective T-cell subsets.

In this study, using a skin-blister induction model we established for dengue(*7*), we investigated the phenotypic, functional and transcriptional signatures of T-cells in matched skin and blood samples from 73 adult dengue patients and 10 healthy volunteers. We found that CD4^+^ and CD8^+^ T-cell responses to DENV infection were highly enriched in the skin compared to the matched blood compartment. Activated CD8^+^ T-cells in the skin displayed phenotypic and transcriptional features of T_RM_ cells including expression of the core gene signature of human T_RM_ cells(*18*), supporting their differentiation into T_RM_ cells. The magnitude of the CD8^+^ T-cell response in the skin positively correlated with that present in the blood of the same patients, and both responses associated with protection from hospitalization in this cohort. These data provide novel insights into the biology and differentiation kinetics of T_RM_ cells in a human virus infection, and support a protective role of skin-resident and circulating CD8^+^ T-cells in dengue, with implications for vaccine design.

## Results

### Activated and proliferating T-cells are enriched in the skin in dengue

To investigate T-cell responses in matched skin and blood compartments during DENV infection we recruited 83 adult study participants, including 73 patients with confirmed dengue (40 ± 13 years; mean age ± SD) and 10 healthy volunteers with no serological evidence of prior DENV infection (42 ± 17 years; mean age ± SD; participant details in **Table I**). Study participants donated blood and skin samples for research purposes at two timepoints (**Fig. 1**): Visit 1 timepoint (days 3-5 from fever onset), was chosen to differentiate primary from secondary DENV infection based on DENV IgG titres, as described previously(27) and for identification of the DENV serotype of infection; visit 2 timepoint (days 7-10 from fever onset), which captured the peak of the T-cell response in dengue(*19, 20*), was chosen for blood and skin T-cell analyses (**Fig. 1**). Skin T-cells were obtained from skin suction blisters induced using a clinical grade pump as described(*7, 21*) (**Fig. 2A)**. On the following day, the fluid accumulated in the skin blister was collected and centrifuged to separate cell pellet from supernatant, both used for downstream analyses. The immune cells accumulating in the skin blister are derived from the underlying skin tissue as confirmed by their high expression of the skin-homing marker CLA compared to matched blood T-cells **(Fig. 2B)**. These data confirmed that the utilised skin suction blister method allows isolation of T-cells from the skin with minimal contamination of blood T-cells.

**Fig. 1.**
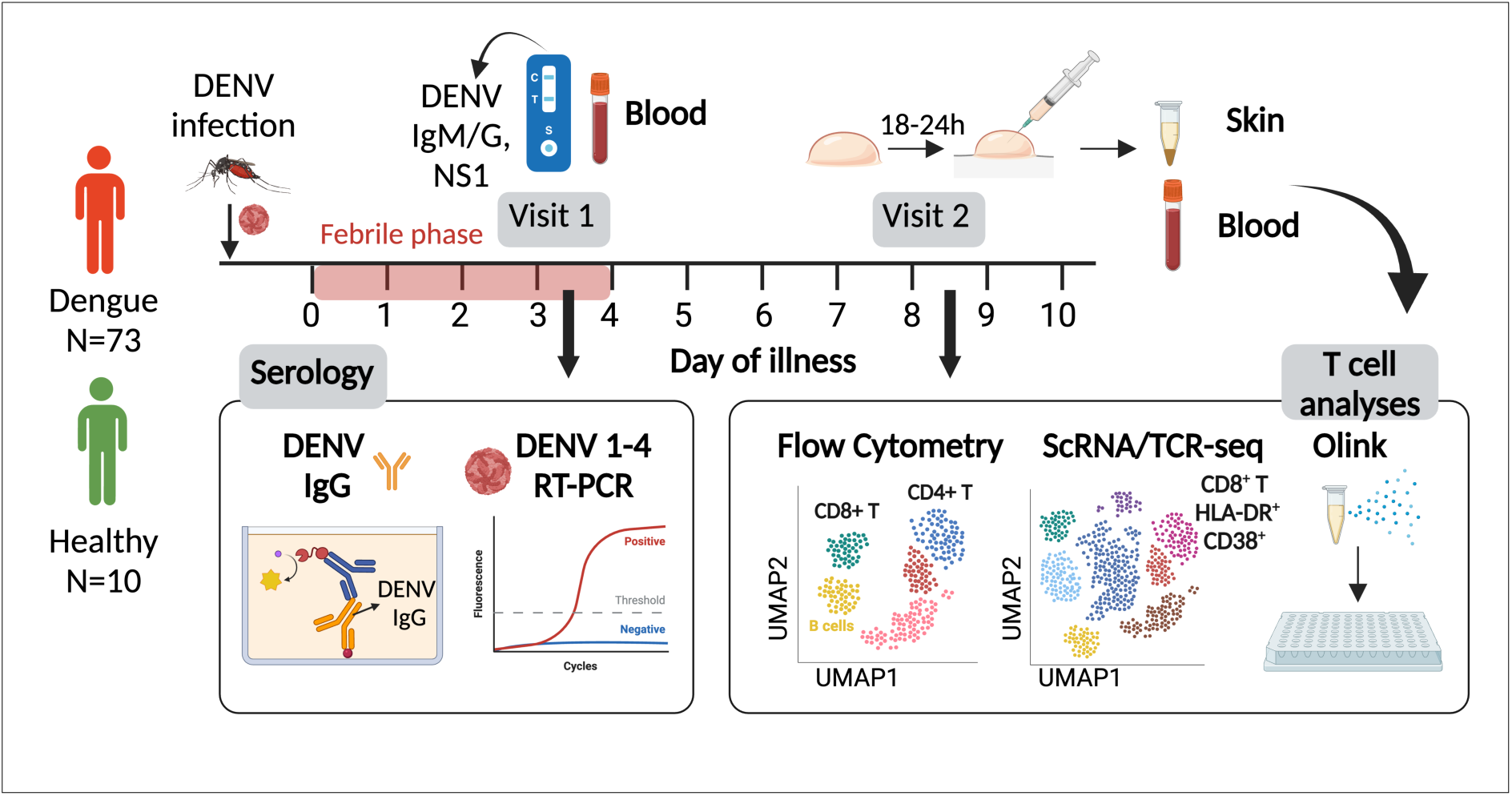
Study design and summary of investigations performed in this study. Day of illness is calculated from the day of fever onset.

**Fig. 2.**
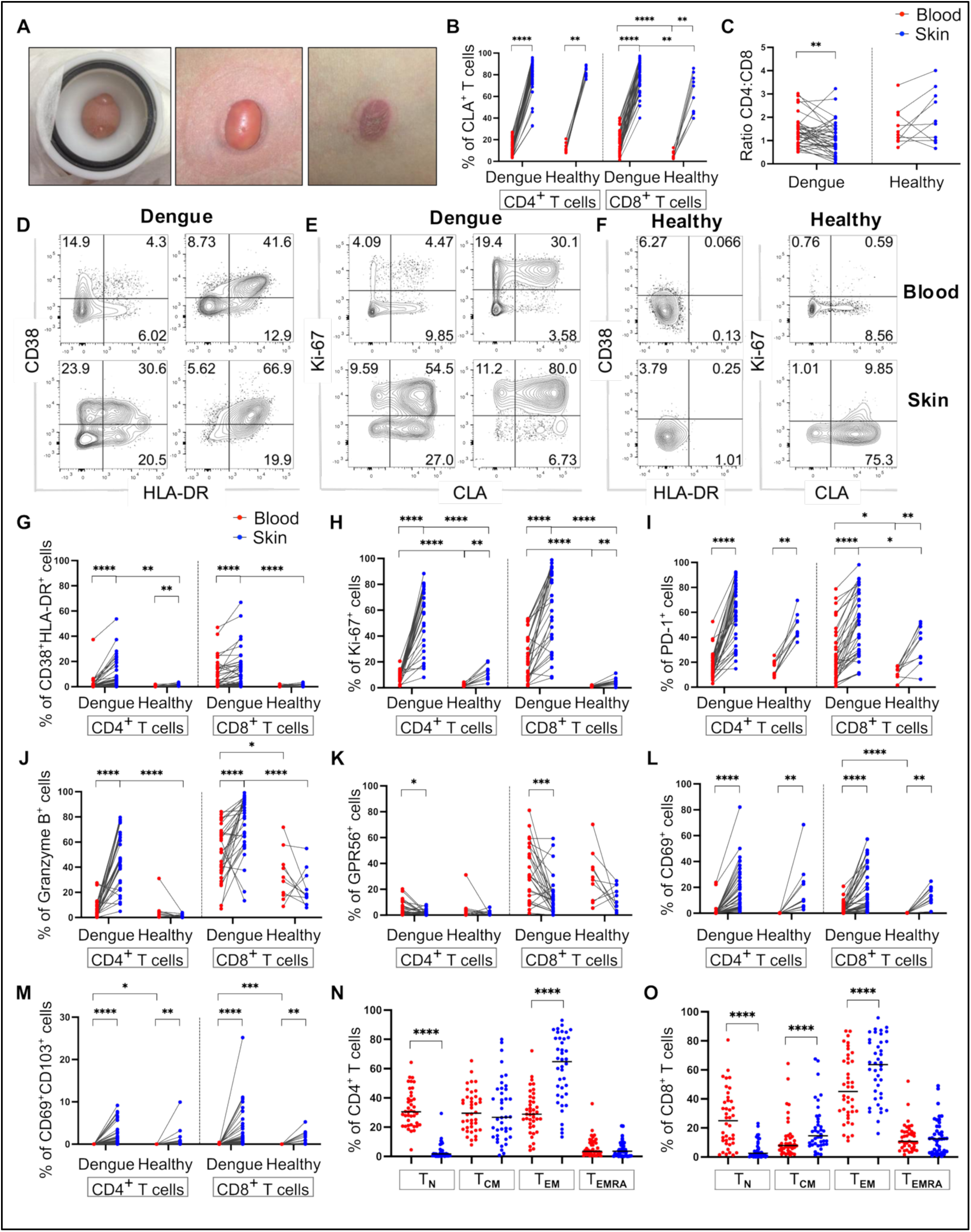
Activated and proliferating T-cells are enriched in the skin of dengue patients. Data from 98 matched blood and skin samples from N=39 dengue patients and N=10 healthy volunteers. **(A)** Photograph of a representative skin blister induced on the forearm of a study participant using a clinical suction pump and chamber, 1 hour (left) and 2 hours (middle) from the start of the procedure, and after aspiration of fluid (right). **(B)** The percentages of blood and skin CLA^+^ CD4^+^ and CD8^+^ T-cells are summarized for patients and healthy volunteers. **(C)** CD4^+^:CD8^+^ T cell ratios in the blood and skin of dengue patients and healthy volunteers. **(D-F)** Representative flow cytometry plots of HLA-DR, CD38 and Ki-67 expression in blood and skin CD4^+^ and CD8^+^ T-cells from a dengue patient (D and E) and healthy volunteer (F, shown for CD4^+^ T-cells). **(G-M)** Frequencies of HLA-DR^+^ CD38^+^ (G) , Ki-67^+^ (H) , PD-1^+^ (I) granzyme B^+^ (J), GPR56^+^ (K), CD69^+^ (L) and CD69^+^ CD103^+^ (M) CD4^+^ and CD8^+^ T-cells expressing in skin and blood of dengue patients and healthy volunteers. (**N-O)** Frequencies of blood and skin T-cell subsets defined by CCR7 and CD45RA expression: T_N_ (naïve): CCR7^+^CD45RA^+^; T_CM_ (T central memory): CCR7^+^CD45RA^−^; T_EM_ (T effector memory): CCR7^−^ CD45RA^−^; T_EMRA_ (T effector memory re-expressing CD45RA): CCR7^−^CD45RA^+^. Datapoints for blood and skin samples for each participant are shown respectively in red and blue. Statistics were determined by Wilcoxon matched-pairs sign rank test between blood and skin, and Mann Whitney t-test between dengue patients and healthy volunteers. For all Figs: *p ≤0.05, **p ≤0.01, ***p ≤0.005, ****p ≤0.0001.

**Table 1:** Clinical information for the dengue patients and healthy volunteers included in the study.

|  | Dengue patients (N = 73) | Healthy donors (N = 10) |
| --- | --- | --- |
| **Age** |  |  |
| Mean ± SD (years) | 40 ± 13 | 42 ± 17 |
| **Sex** |  |  |
| Male | 59 (81%) | 7 (70%) |
| Female | 14 (19%) | 3 (30%) |
| **2009 WHO Classification** |  |  |
| Severe Dengue | 3 (4%) | N.A. |
| Dengue With Warning Signs | 43 (59%) |  |
| Dengue Without Warning Signs | 27 (37%) |  |
| **Serostatus** |  |  |
| Primary infection | 40 (55%) | N.A. |
| Secondary infection | 33 (45%) |  |
| **Serotype** |  |  |
| DENV1 | 2 (3%) | N.A. |
| DENV2 | 34 (47%) |  |
| DENV3 | 18 (25%) |  |
| DENV4 | 1 (1%) |  |
| Unknown (< LOD) | 18 (25%) |  |
| **Hospitalisation** |  |  |
| Outpatient | 22 (30%) | N.A. |
| Inpatient | 31 (42%) |  |
| Outpatient/Inpatient * | 20 (27%) |  |
| **Thrombocytopenia** |  |  |
| With | 53 (73%) | N.A. |
| Without | 20 (27%) |  |
| **Rash** |  |  |
| With | 27 (37%) | N.A. |
| Without | 45 (62%) |  |
| **Highest hematocrit** |  |  |
| Mean ± SD (%) | 45.7 ± 4.0 | Not recorded |
| **Lowest platelet count** |  |  |
| Mean ± SD ($\times 10^3/\mu\text{L}$) | 73.2 ± 51.8 | Not recorded |

We first investigated CD4^+^ and CD8^+^ T-cell activation and proliferation within the skin and blood compartments of 49 participants (39 dengue patients and 10 healthy volunteers) and asked whether skin T-cells possessed distinct phenotypic and functional features compared to their circulating counterparts. Peripheral blood mononuclear cells (PBMCs) and cells from the skin blister aspirates, defined hereinafter as “blood” and “skin” cells, were analysed by flow cytometry for expression of CD4^+^ and CD8^+^ T-cell markers of activation and proliferation (HLA-DR, CD38, CD69, PD-1, Ki-67), tissue residence (CD69, CD103, CLA), differentiation (CCR7, CD45RA) and cytotoxicity (granzyme B, GPR56). In contrast to the healthy volunteers, dengue patients displayed higher CD4^+^:CD8^+^ T-cell ratios in blood compared to skin, consistent with an expansion or influx of CD8^+^ T-cells into the skin compartment during DENV infection **(Fig. 2C)**. Furthermore, CD4^+^ and CD8^+^ T-cells in the skin of dengue patients displayed increased activation and proliferation compared to their blood counterparts **(Fig. 2D-E; 2G-I)**. Low levels of activated and proliferating T-cells could be detected in the skin but not in the blood of healthy volunteers, suggesting that T-cells in the skin may undergo turnover in steady state **(Fig. 2G-I)**. The frequencies of PD-1^+^ T-cells were also significantly higher in the skin compared to the blood of healthy volunteers, in line with previous reports(*22*) **(Fig. 2I)**. In dengue patients but not in healthy volunteers, skin CD4^+^ and CD8^+^ T-cells contained higher frequencies of granzyme B^+^ cells compared to their blood counterparts **(Fig. 2J)**. GPR56, a G protein-coupled receptor described to mark blood cytotoxic T-cells (*23, 24*) was expressed at higher levels in blood compared to skin CD4^+^ and CD8^+^ T-cells of dengue patients, while expression was similar across these two compartments in T-cells from healthy volunteers **(Fig. 2K)**. Skin CD4^+^ and CD8^+^ T-cells from both dengue patients and healthy volunteers expressed significantly higher levels of CD69 compared to their blood counterparts, with a proportion of skin CD69^+^ cells co-expressing CD103 **(Fig. 2L-M)**. CD69^+^ and CD69^+^ CD103^+^ CD8^+^ T-cells were increased in the blood of dengue patients compared to that of healthy volunteers, although their levels were lower compared to those of their skin counterparts **(Fig. 2L-M)**. In dengue patients, skin CD4^+^ and CD8^+^ T-cells were mainly CCR7^−^CD45RA^−^ (T effector memory; T_EM_) followed by CCR7^+^CD45RA^−^ (T central memory cells; T_CM_). In contrast, blood contained CCR7^+^CD45RA^+^ (naïve; T_N_), T_CM_ and T_EM_ CD4^+^ and CD8^+^ T-cells and CD45RA^+^ (T effector memory RA re-expressing; T_EMRA_) CD8^+^ T-cells **(Figure 2N, O)**. Similarly, CD4^+^ and CD8^+^ T-cells in the skin of healthy volunteers comprised respectively of T_EM_ and T_CM_ cells, as well as T_EMRA_ cells for CD8^+^ T-cells (**Fig. S1A-B**). In summary, our data demonstrate that activated and proliferating T-cells expressing high levels of granzyme B were highly enriched in the skin compared to the blood of dengue patients, suggesting that the skin is a key site for T-cell immunosurveillance of DENV during acute infection.

### Dramatic reshaping of the skin T-cell compartment in dengue

To investigate the phenotypic and functional features of skin and blood T-cells, we performed unsupervised analyses using dimensionality reduction algorithms. To this end, we selected flow cytometry standard (FCS) files from 54 matched skin and blood samples derived from 17 dengue patients and 10 healthy volunteers, which had been analysed using identical flow cytometer settings. These files were downsampled to the same number of cells, concatenated and analysed using uniform manifold approximation and projection (UMAP) and Phenograph cluster analysis to identify T-cell populations enriched within blood or skin compartments. Our findings show that blood and skin CD4^+^ and CD8^+^ T-cells mapped to distinct areas of the UMAP plots in both dengue patients and healthy volunteers (**Fig. 3A**). A partial overlap of skin and blood CLA^+^ Ki-67^+^ T-cells was observed in dengue patients (**Fig. 3A, B**), suggesting that these cells recirculate between the two compartments during infection. Phenograph analyses identified 14 different CD3^+^ T-cell clusters, which included 5 CD4^+^ T-cell and 7 of CD8^+^ T-cell clusters (respectively clusters 1, 3, 4, 7, 9 and 2, 5, 6, 8, 10, 11, 14; **Fig. 3C**) as well as 2 clusters with low CD4 and CD8 expression (clusters 12 and 13; **Fig. 3C**). The frequencies of T-cells within these clusters and the relative mean fluorescence intensity (MFI) expression of the markers analysed are summarized in a heatmap (**Fig. 3D**). These 14 clusters largely distinguished blood T-cells from their skin counterparts, albeit with some clusters being shared between the two compartments. Skin T-cells from healthy volunteers mapped predominantly within clusters 1 and 5, followed by cluster 10 and were largely non-overlapping with their blood counterparts (**Fig. 3E**). In contrast, skin T-cells from dengue patients largely localized within clusters 2 and 9, which comprise respectively of CD8^+^ and CD4^+^ T-cells that are highly activated and proliferating, display cytotoxic capacity (granzyme B^high^), and high expression of CLA and CD69, but low expression of CD103 and CCR7 (**Fig. 3D**). These cells were enriched in the skin of dengue patients, were present albeit at lower frequencies in their blood counterparts and were largely absent in healthy volunteers (**Fig. 3E**). The phenotypic/functional features of these cells (CLA^+^, PD-1^+^, Ki-67^high^, CD69^+^, granzyme B^high^) strongly resembled those of DENV-specific T-cells detected in the blood of dengue patients using DENV peptide-HLA class-I tetramers at day 7-10 of illness(*24*), which were shown to be distinct to that of “bystander activated” T-cells in dengue that lack expression of CLA and are specific for antigens unrelated to DENV(*7*). Based on these considerations, these cells may represent *bona fide* DENV-specific T-cells and are herein named “responding T-cells”.

**Fig. 3.**
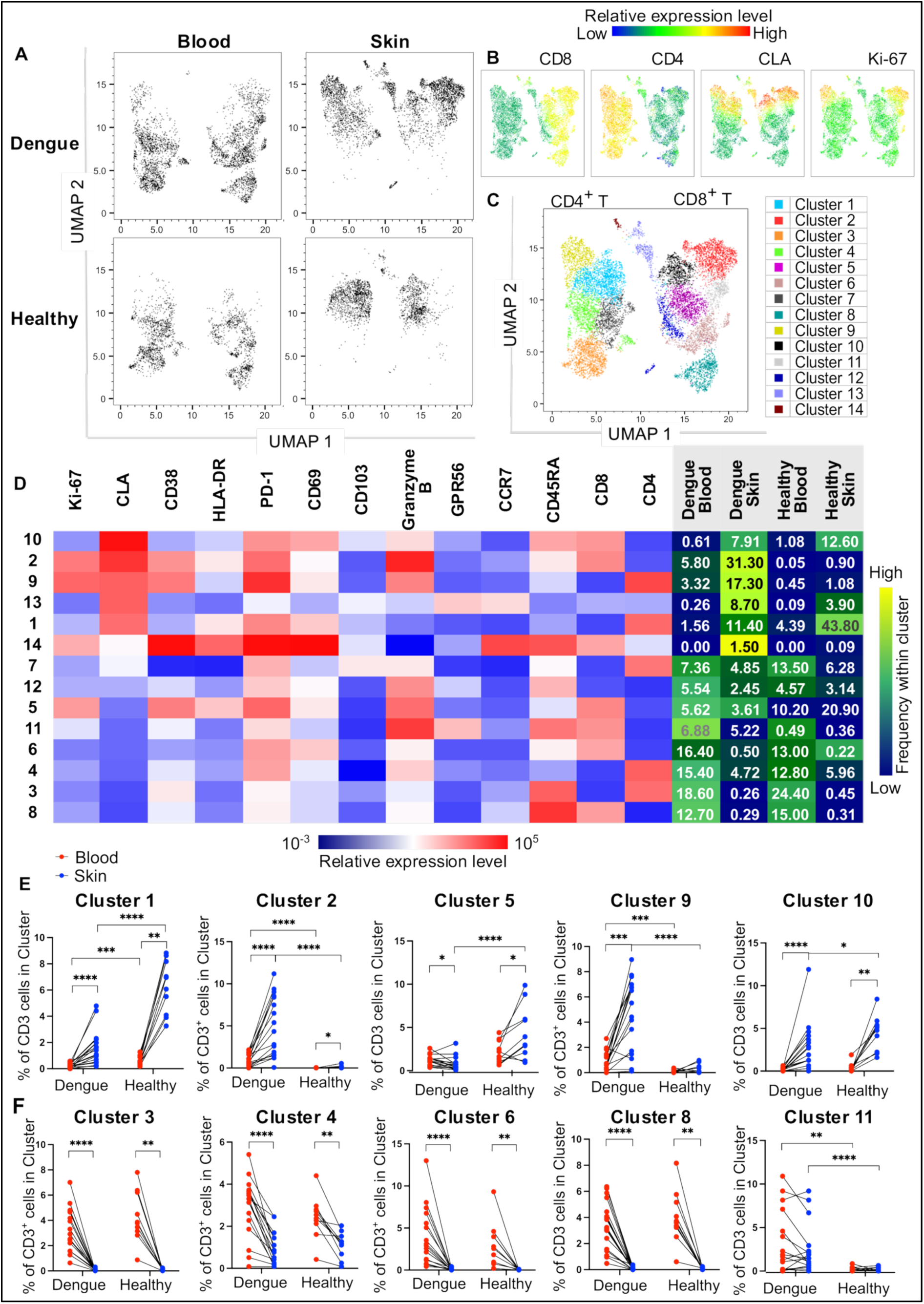
Skin and blood T-cells are largely distinct. Data from 54 matched skin and blood samples from N=17 dengue patients and N=10 healthy volunteers**. (A-B)** UMAP plots of blood and skin CD3^+^ T-cells from dengue patients and healthy volunteers. Expression of CD8, CD4, CLA and Ki-67 within the UMAP is shown in B. **(C)** The 14 T-cell clusters identified by Phenograph within CD3^+^ cells are shown. Each colour corresponds to a cluster. **(D)** Relative mean fluorescence intensities (MFI) of the analysed markers within each the cluster are shown in the heatmap. The mean frequencies of T-cells within each cluster in blood and skin of dengue patients and healthy volunteers are shown in the columns (right; sum of the mean frequencies is 100% within each column). **(E, F)** Frequencies of cells within clusters enriched in the skin (E) or blood (F) of dengue patients and healthy volunteers. Statistics were calculated by Wilcoxon matched-pairs sign rank test between blood and skin, and Mann-Whitney t-test between healthy volunteers and patients.

Dengue patients’ blood T-cells mapped predominately within clusters 3, 4, 6 and 8 followed by cluster 11 (**Fig. 3D, F**). The two most abundant clusters of CD4^+^ T-cells (clusters 3 and 4) and CD8^+^ T-cells (clusters 6 and 8) in the blood of dengue patients displayed lower expression of HLA-DR, CD38, granzyme B, Ki-67 and CLA compared to T-cells enriched in the skin. Cluster 11 containing PD-1^+^ CD8^+^ CD45RA^+^ CCR7^−^ (T_EMRA_ cells) with high cytotoxic potential (granzyme B^+^ GPR56^+^) was present in the blood and skin of dengue patients but largely absent in both compartments of healthy volunteers (**Fig. 3F**). In contrast, clusters 1 and 10 containing respectively CD4^+^ and CD8^+^ T-cells expressing CLA, PD-1 and CD69 but lacking expression of activation and proliferating markers were present in the skin of both healthy volunteers and dengue patients, although frequencies were higher in the former (**Fig. 3D, E**).

These cells may be resting skin resident T-cells. Overall, skin T-cells in dengue patients differed substantially from those in healthy volunteers, while blood T-cells looked more similar between dengue patients and healthy volunteers.

In summary, unsupervised analyses show that blood and skin T-cells were largely phenotypically distinct in steady state and during DENV infection. In dengue patients, skin T-cells were dramatically distinct from both their blood counterparts and skin T-cells of healthy volunteers, suggesting that the skin compartment undergoes extensive reshaping during DENV infection.

### Skin and blood DENV-specific CD8^+^ T-cells display similar features

As blood and skin T-cells appeared to be largely distinct populations of cells, we next asked whether the features of DENV-specific CD8^+^ T-cells are distinct in these two compartments. DENV-specific T-cells were analysed using peptide-HLA class I pentamers directly *ex vivo*. We previously showed that DENV NS3 and NS5 are the major targets of the CD8^+^ T cell response(*20*), with NS3_1608-1617_ and NS5_2610-2618_ epitopes being the most frequently recognised epitopes restricted to HLA-A*11:01, a commonly expressed HLA-type in the Singapore population(*24, 25*). Here, we identified 8 HLA-A*1101^+^ dengue patients within our cohort whose CD8^+^ T-cells responded with detectable frequencies to at least one of these epitopes, and analysed their blood and skin cells by flow cytometry using HLA-A*11:01-NS3_1608-1617_ and NS5_2610-2618_ pentamers matched to the DENV serotype of infection for each patient. To evaluate the features of the DENV-specific T-cells, cells were co-stained with HLA-A*11:01 pentamers and antibodies targeting phenotypic and functional T-cell markers. The frequencies of CD8^+^ HLA-A*11:01-NS3_1608-1617_/NS5_2610-2618_ pentamer^+^ cells varied between patients but were consistently higher in the skin compared to the blood in 6 out of 8 patients, supporting our above findings of a more robust T-cell response in the skin (**Fig. 4 A, B**). For 2 of these patients, we obtained a sufficient number of events to allow a more in-depth UMAP and Phenograph cluster analyses of pentamer^+^ skin versus blood CD8^+^ T-cells. Interestingly, in both patients, the manually gated pentamer^+^ cells mapped to a single overlapping cluster of the UMAP and Phenograph plots (**Fig. 4C, D**: cluster in green/cluster 2 and **Fig. S3).** The proportions of pentamer^+^ skin CD8^+^ T-cells were on average approximately 10-fold higher compared to those in the blood (**Fig. 4E**: cluster 2, 25.1% versus 2.24%; **Fig. S3**). Cells in cluster 2 displayed a phenotype which is characteristic of DENV-specific CD8^+^ T-cells, namely CD45RA^+^ CCR7^low^ and with high expression of Ki-67, CLA, CD69, CD38, HLA-DR, PD-1 and granzyme B (**Fig. 4E**).

**Fig. 4:**
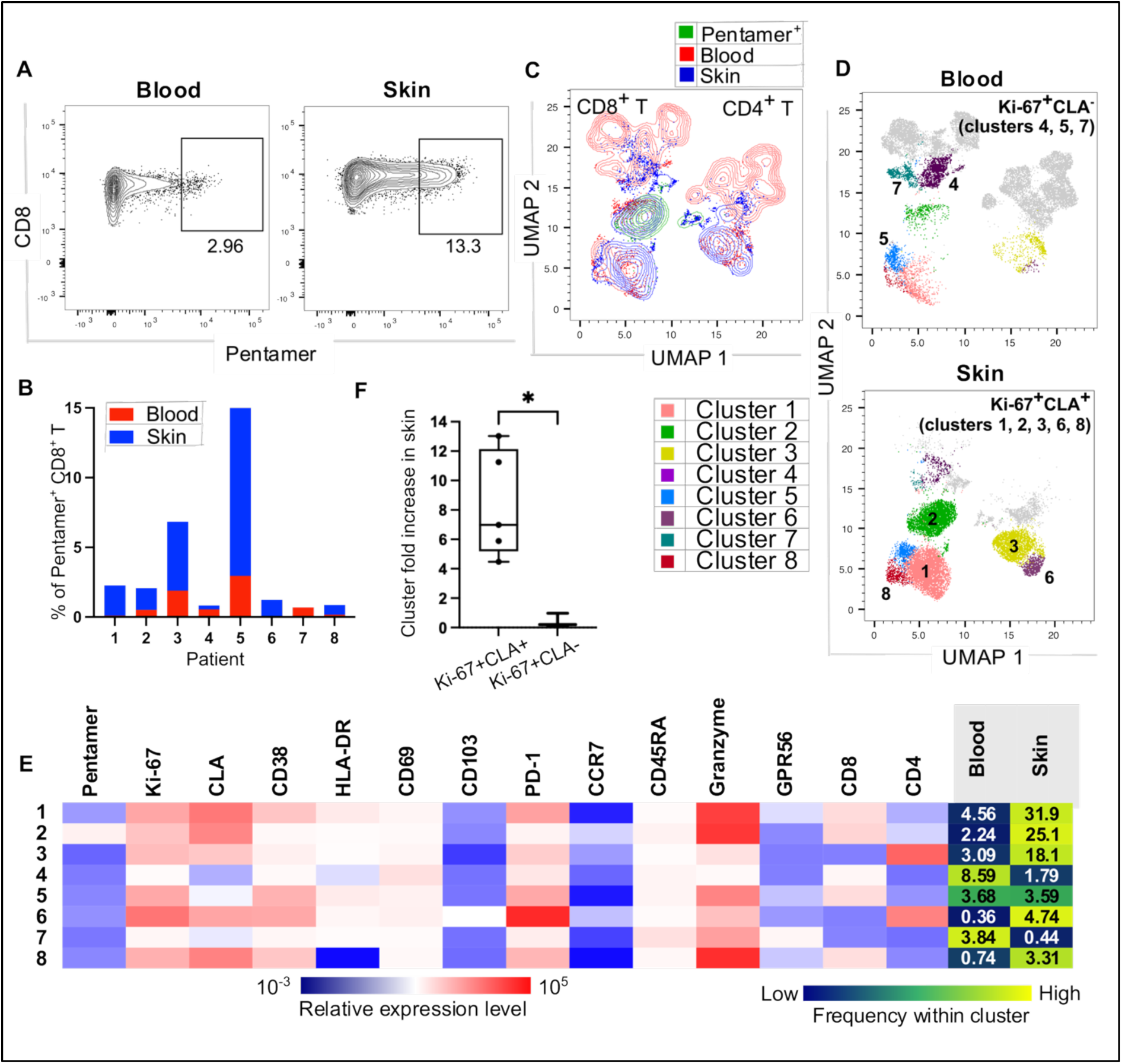
DENV pentamer^+^ and Ki-67^+^ CLA^+^ T-cells are enriched in the skin. **(A)** Representative flow cytometry plots showing staining profiles for pooled DENV NS3_1608-1617_-HLA-A*11:01 and NS5_2610-2618_-HLA-A*11:01 pentamers (defined herein DENV pentamers) of CD3^+^ T-cells in matched blood and skin of a dengue patient. Pentamers were matched to the DENV serotype of infection **(B)** Frequencies of DENV pentamer^+^ CD8^+^ T-cells in 16 matched blood and skin samples from N=8 dengue patients. **(C)** UMAP plots showing the localization of manually-gated DENV pentamer^+^ CD8^+^ T-cells (green) within blood (red) and skin (blue) CD3^+^ T cells in 4 matched blood and skin samples from N=2 patients. **(D)** Shown are the clusters identified by Phenograph that correspond to Ki-67^+^CLA^+^ and Ki-67^+^CLA^−^ CD4^+^ and CD8^+^ T-cells shown for blood and skin cells. **(E)** Heatmap showing MFI expression of the analysed markers in Ki-67^+^CLA^+^ and Ki-67^+^CLA^−^ clusters from D. The mean frequencies of each cluster are shown within blood and skin T cells. **(F)** Shown is the fold increase of Ki-67^+^CLA^+^ and Ki-67^+^CLA^−^ T-cells in skin versus blood. Statistics in E were calculated using Mann Whitney t-test.

Our previous work using DENV peptide-HLA class-I tetramers showed that CLA and Ki-67 can be used to identify *bona fide* DENV-specific T-cells that are responding to DENV(*24*). To analyse the broader pool of responding T-cells, we therefore analysed the features of Ki-67^+^ CLA^+^ CD4^+^ and Ki-67^+^ CLA^+^ CD8^+^ T-cells (**Fig. 4D**). Ki-67^+^ CLA^+^ CD8^+^ T-cells mapped to 3 distinct clusters, cluster 2 which contained the pentamer^+^ cells and clusters 1 and 8 which were adjacently located and contained CD8^+^ T-cells expressing similar phenotypic and functional markers, except for HLA-DR which showed decreased expression in cluster 8 (**Fig. 4D, E**). Ki-67^+^ CLA^+^ CD4^+^ T-cells mapped to 2 adjacently located clusters (clusters 3 and 6). All 5 clusters of Ki-67^+^ CLA^+^ T-cells were more abundant in the skin compared to matched blood (**Fig. 4F**). We identified 3 clusters of cells which were Ki-67^+^ but did not express CLA, these were CD8^+^ T-cells (clusters 4 and 5) or CD4^low^CD8^low^ T-cells (cluster 7) and were enriched in blood compared to skin (clusters 4 and 7) or present at similar frequencies in these two compartments (cluster 5; **Fig. 4D, F)**.

In summary, we show that DENV-specific pentamer^+^ CD8^+^ T-cells and Ki-67^+^ CLA^+^ CD4^+^/CD8^+^T-cells were highly enriched in the skin compared to blood. Skin and blood pentamer^+^ CD8^+^ T-cells were located within the same cluster of cells, suggesting similar expression of the analysed markers in DENV-specific skin and blood T-cells and potentially reflecting recent migration of these cells from skin to blood.

### Increased CD8^+^ Ki-67^+^ CLA^+^ T cell responses in non-hospitalized dengue patients

We further investigated the relationship between skin and blood responding Ki-67^+^ CLA^+^ T-cells in a larger number of patient samples and asked whether the frequencies of these cells in skin and/or blood correlate with dengue disease outcomes. To achieve this, we compared the frequencies and phenotypic/functional features of Ki-67^+^ CLA^+^ CD4^+^ and Ki-67^+^ CLA^+^ CD8^+^ T-cells in 78 matched skin and blood samples from 29 dengue patients and 10 healthy volunteers. The frequencies of Ki-67^+^ CLA^+^ T-cells were strikingly higher in dengue patients compared to healthy volunteers, and in skin compared to blood (**Fig. 5A**). In dengue patients, Ki-67^+^CLA^+^ T-cells showed similar co-expression of HLA-DR and CD38 (**Fig. 5B**). However, the frequencies of cells expressing PD-1, granzyme B and the skin residency markers CD69/CD103 were significantly higher in skin versus blood Ki-67^+^ CLA^+^ CD4^+^ and CD8^+^ T-cells (**Fig. 5B-G**). Skin and blood Ki-67^+^ CLA^+^ CD4^+^ and CD8^+^ T-cells were predominantly T_CM_ and T_EM_ cells, although skin Ki-67^+^ CLA^+^ T-cells contained a minor fraction of cells that were phenotypically naïve and T_EMRA_ cells (**Fig. 5H-I**). Interestingly, the frequencies of skin Ki-67^+^ CLA^+^ CD4^+^ T-cells varied largely amongst patients and did not correlate with those present in matched blood, suggesting that generation of these cells may be independently controlled (**Fig. 5J**). In contrast, the frequencies of Ki-67^+^ CLA^+^ CD8^+^ T in the skin directly correlated with those present in the blood, suggesting there may be recirculation of these cells between the two compartments (**Fig. 5K**). To assess the correlation of T-cell responses with clinical outcomes we stratified patients based on whether they required hospitalization (inpatients) or were deemed fit to return to their home (outpatients), following a clinical assessment by expert clinicians. The frequencies of skin Ki-67^+^ CLA^+^ CD4^+^ T-cells were highest in patients with primary infection and showed a higher trend in outpatients compared to inpatients, while no differences were observed for blood Ki-67^+^ CLA^+^ CD4^+^ T-cells (**Figure 5L-M; 5P-Q)**. In contrast, outpatients displayed higher frequencies of both blood and skin Ki-67^+^ CLA^+^ CD8^+^ T-cells compared to inpatients. Patients with primary DENV infection displayed increased trends of Ki-67^+^ CLA^+^ CD8^+^ T-cells although these differences were not statistically significant (**Fig. 5N-O; 5R-S**).

**Fig. 5:**
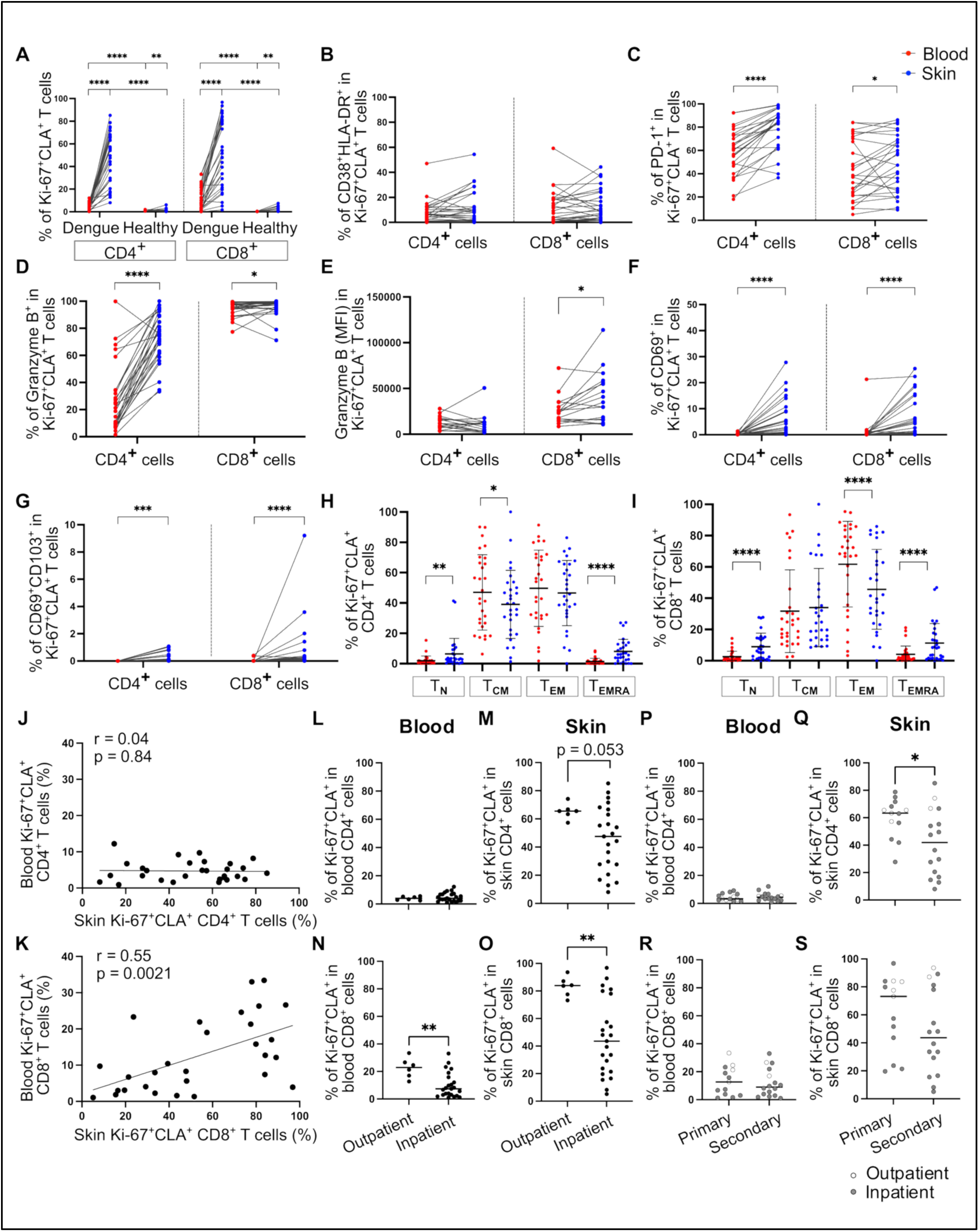
The frequencies of skin Ki-67^+^ CLA^+^ CD8^+^ T-cells are directly correlated and associate with improved disease outcomes. Data from 78 matched skin and blood samples from N=29 dengue patients and N=10 healthy volunteers. **(A)** Frequencies of Ki-67^+^ CLA^+^ cells in blood and skin CD4^+^ and CD8^+^ T-cells. **(B-G)** Frequencies of Ki-67^+^ CLA^+^ CD4^+^ and Ki-67^+^ CLA^+^ CD8^+^ T-cells expressing the indicated markers of activation proliferation, tissue-residence & cytotoxicity in the blood and skin. MFI values are also shown for granzyme B in (E). **(H-I)** Differentiation status of Ki-67^+^ CLA^+^ CD4^+^ and CD8^+^ T-cells defined by expression of CCR7 and CD45RA. **(J-K)** Correlation of blood and skin Ki-67^+^ CLA^+^ CD4^+^ (J) and Ki-67^+^ CLA^+^ CD8^+^ T-cells (K) using Spearman’s rank correlation test. **(L-O).** Frequencies of Ki-67^+^ CLA^+^ CD4^+^ (L-M) and CD8^+^ T-cells (N-O) are shown in patients stratified by patient hospitalization status. **(P-S).** Frequencies of Ki-67^+^ CLA^+^ CD4^+^ (P-Q) and CD8^+^ T-cells (R-S) are shown in patients stratified by primary/secondary infection. Statistics were calculated using Wilcoxon matched-pairs sign rank test between blood and skin, and Mann-Whitney t-test between healthy volunteers and patients (A-I) and inpatients and outpatients (L-S).

In summary, we show that responding Ki-67^+^ CLA^+^ CD4^+^ and CD8^+^ T-cells were enriched in the skin compartment. The frequencies of Ki-67^+^ CLA^+^ CD8^+^ T-cells in skin and blood were directly correlated and were increased in patients that did not require hospitalization compared to those that did, suggesting a protective role of CD8^+^ T-cells.

### Increased T-cell cytokine responses in non-hospitalized patients

To confirm whether there was a superior skin T-cell response in outpatients compared to inpatients on a larger number of patient samples, we analysed skin blister fluid supernatants from 69 patients, including 22 outpatients, 18 outpatients/inpatients and 29 inpatients, for the presence of 45 cytokines/chemokines using the Olink® Target 48 Cytokine panel. Outpatients/inpatients were outpatients at Visit 1 but were later hospitalised due to worsening conditions. Outpatients displayed higher levels of Interleukin (IL)-2, IL-17A, IL-17F and IL-17C in the skin compared to inpatients, with a stepwise decrease of these cytokines from outpatients to outpatients/inpatients and inpatients (**Fig. 6A-D**). IL-2 and IL-17 cytokines are part of the core transcriptional signature described for CD4^+^ and CD8^+^ T_RM_ cells(*18*). The levels of IL-27, a member of the IL-12 cytokine family, which supports T-cell differentiation was also higher in outpatients compared to outpatients/inpatients and inpatients (**Fig. 6E**). Outpatients also displayed higher levels of hepatocyte growth factor (HGF), tumour necrosis factor superfamily member 12 (TNFSF12) and CCL-19, a chemokine which mediates recruitment of CCR7^+^ T-cells (**Fig. 6F-H**). In contrast, chemokines associated with the recruitment of cell-types including leukocytes, neutrophils and eosinophils (respectively CCL7, CXCL8 and CCL11) were increased in outpatients/inpatients compared to outpatients, although differences were less striking (**Fig. 6I-K**).

**Fig. 6.**
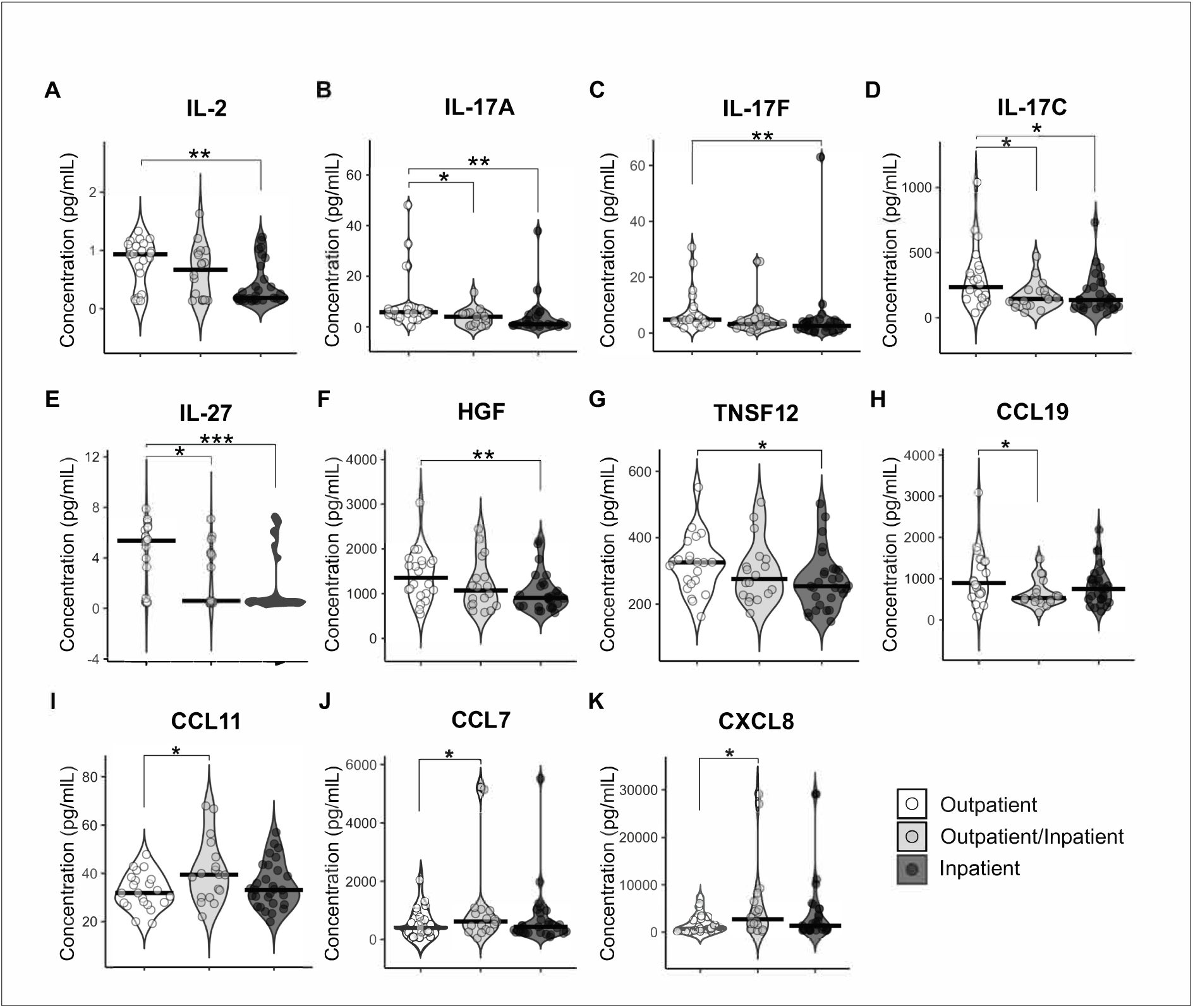
Increased T-cell related cytokines in the skin correlate with improved clinical outcomes. Cytokines were measured using Olink Target 48 proteomics in skin blister fluids of N=69 dengue patients. Shown are the concentrations (pg/ml) of IL-2 (A), IL-17A (B), IL-17F (C), IL-17C (D), IL-27 (E), HGF (F), TNSF12 (G), CCL19 (H), CCL11 (I), CCL7 (J) and CXCL8 (K). Data is stratified by patients’ hospitalization status (Outpatient, N=22; Outpatient/Inpatient, N=18; Inpatient, N=29). Statistics were calculated by One-way ANOVA.

In summary, our data shows increased levels of T-cell related cytokines including IL-2, IL-17 and IL-27 in the skin of outpatients compared to inpatients, supporting the association of a superior skin T-cell response with improved clinical outcomes in this dengue cohort.

### Activated skin CD8^+^ T-cells display features of T_RM_ cells

To gain a more in-depth understanding of the features and clonal relationship of skin and blood T-cell responses in dengue, we performed paired gene expression and T-cell receptor (TCR) analyses by 10X Genomics single-cell (sc) RNA-sequencing (RNA-seq) and TCR-sequencing (TCR-seq) of matched skin and blood activated (HLA-DR^+^ CD38^+^) CD8^+^ T-cells isolated by cell sorting (days 7-10 from fever onset; patient details in **Table S1** and **Table 1**). ScRNA/scTCR-seq was performed on a combined total of 8,175 single-cells derived from 6 matched skin and blood of 3 dengue patients. UMAP analyses of the gene expression data identified clusters of T-cells that were clearly distinct between blood and skin, and a cluster of proliferating cells (cycling) that contained both skin and blood CD8^+^ T-cells (**Fig. 7A**). Skin CD8^+^ T-cells displayed increased expression of genes encoding for granzyme B (*GZMB)*, granulysin (*GNLY*) and perforin (*PRF1),* which are critical for cellular cytotoxicity, with granulysin and granzyme B expression progressively increasing in clusters corresponding to blood, blood cycling, skin and skin cycling CD8^+^ T-cells. However, other genes also linked to cytotoxicity displayed increased expression in blood CD8^+^ T-cells compared to their skin counterparts (serglycin, *SRGN*; natural killer cell granule protein 7, *NKG7* and granzyme H, *GZMH*) (**Fig. 7B**). Skin CD8^+^ T-cells displayed increased expression of the skin and peripheral tissue residency markers CD69, CD103 (*ITGAE*), CD49 (*ITGA1*), CXCR6 as well as PD-1 (*PDCD1*), compared to their blood counterparts. Conversely, *EOMES, KLF2, S1PR1* and *TBX21* which encode for transcription factors described to be downregulated in T_RM_ cells (respectively Eomes; Kruppel factor 2, KLF2; Sphingosine-1-phosphate receptor 1, S1PR1 and T-bet) displayed decreased expression in skin compared to blood CD8^+^ T-cells. CX3CR1 expression was also decreased in skin T-cells compared to their blood counterparts (**Fig. 7C**). These data suggest that CD8^+^ T-cells activated during DENV infection differentiate into T_RM_ cells by days 7-10 from illness onset.

**Fig. 7.**
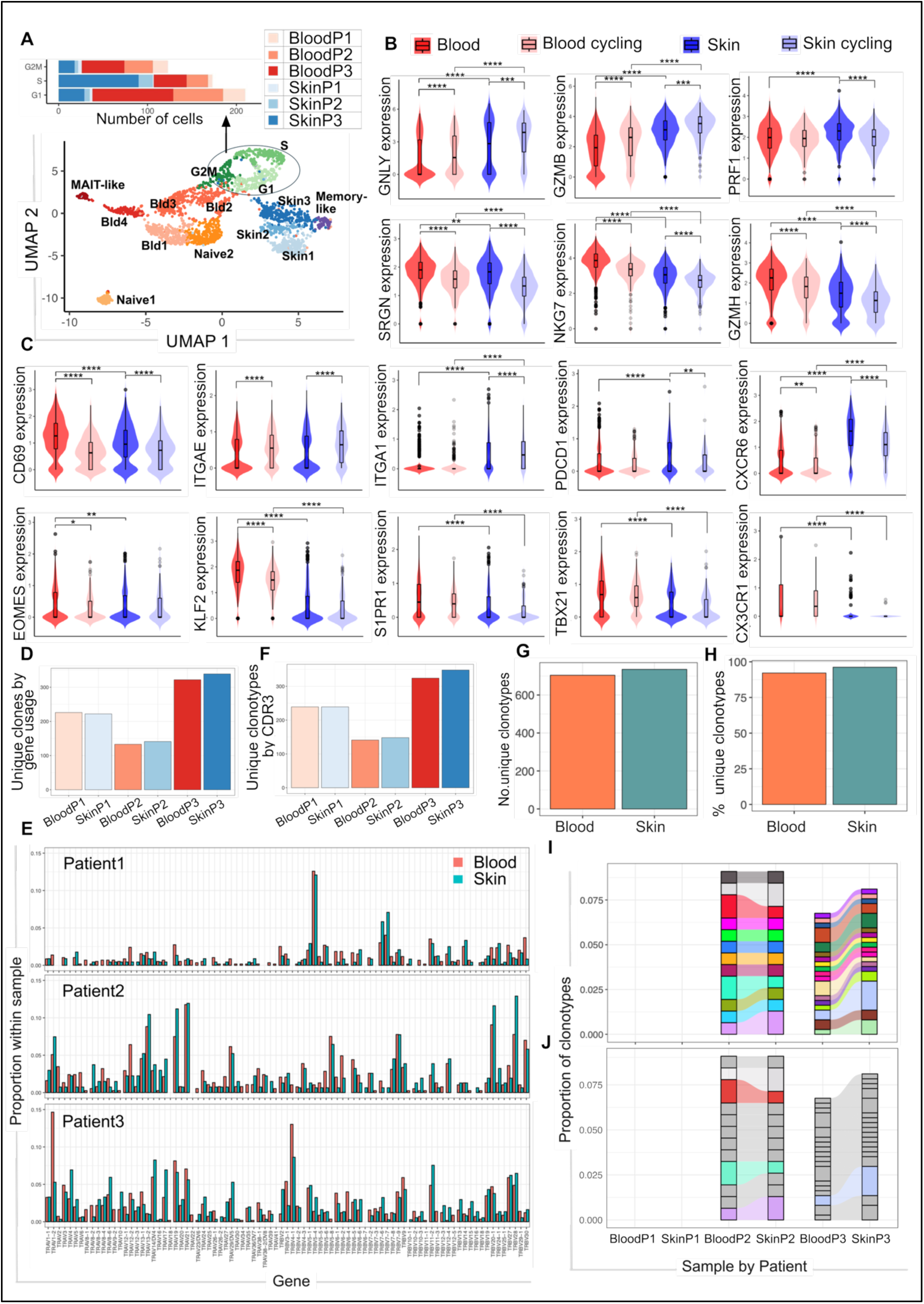
Activated CD8^+^ T-cells in the skin in dengue display features of T_RM_ cells. **(A)** UMAP plot showing the distribution of skin (blue), blood (red-orange) and cycling (green) CD8+ T-cells based on single-cell gene expression. The number of cells from blood and skin of each patient which are located in the different phases of the cell cycle, are indicated in the top graph. **(B)** Gene expression levels of genes involved in cytotoxicity are shown for CD8^+^ T-cells and cycling CD8^+^ T-cells in blood and skin. GNLY: granulysin, GZMB: granzyme B, PRF1: perforin, SRGN: serglycin, NKG7: natural killer cell granule protein 7, GZMH: granzyme H. **(C)** Expression levels are shown for genes known to be upregulated or downregulated in T_RM_ cells (respectively, top and bottom panel) in blood and skin CD8^+^ T-cells and cycling CD8^+^ T-cells. **(D-H)** Number of unique clones defined by TRAV and TRBV gene usage is shown for blood and skin CD8^+^ T-cells in each patient (D) and summarised for all patients (G). Expression of TRAV and TRVB genes in blood and skin activated CD8^+^ T-cells is shown for each patient (E). Number of unique clonotyes defined by CDR3 regions are shown for each patient (F) and percentage of unique clonotypes is summarised for all patients (H), downsampled by each patient’s lower limit of cell counts. **(I)** The proportion of clonotypes, defined by the CDR3 amino acid sequences, shared by skin and blood samples is shown for each patient after downsampling for each patient’s lowest cell count sample. Colours identify unique clonotypes. **(J)** Shared clonotypes that are expanded (proportion > 0.01) in skin and blood of patients 2 and 3 are shown. Statistics in B-C are calculated using One-way Anova with Benjamini-Hochberg correction. P1, P2 and P3 indicates respectively patients 1, 2 and 3.

We next measured the clonal diversity of matched skin and blood CD8^+^ T-cells and their clonotype overlap. To this end, we analysed the expression of TCR α and β chain variable genes (TRAV & TRBV) and the unique complementarity-determining regions (CDR3) that determine the CD8^+^ T-cell clonotypes. For all patients analysed, there was extensive sharing of TRAV and TRBV gene usage between skin and blood CD8^+^ T-cells (**Fig. 7D, E**). Analyses of the unique CDR3-defined clonotypes showed similar number of clonotypes in matched blood and skin CD8^+^ T-cells, with a total of >600 unique clonotypes identified across the 3 patients, comprising of >80% of clonotypes within the activated CD8^+^ T-cell population (**Fig 7F-H**). Analyses of the proportion of clonotype sharing by skin and blood CD8^+^ T-cells revealed 12 and 17 shared clonotypes in patients 2 and 3, respectively (**Fig. 7I**), with 4 of these clonotypes being expanded in size, ranging from small to large/hyperexpanded clonal groups (clonotype proportions ≤0.01 and >0.01, respectively; **Fig. 7J**). These shared CD8^+^ T-cell clonotypes accounted for approximately 7-9% of total clonotypes within activated CD8^+^ T-cells of patients 2 and 3. We did not identify shared clonotypes between skin and blood CD8^+^ T-cells of patient 1 (**Fig. 7I)**.

In summary, these data show that activated and responding skin CD8^+^ T-cells in DENV infection displayed transcriptional features of T_RM_ cells, supporting the differentiation of these cells into long-lived memory T-cells within the skin tissue at the time of infection. Tissue-resident and circulating CD8^+^ T-cells exhibited a moderate clonotypic diversity and some evidence of clonotype sharing in 2 out of 3 patients, supporting the notion that T_RM_ cells can recirculate between tissues and blood during infection. Further studies are needed to define the extent of trafficking of T-cells responding to DENV infection between the blood and skin compartments and potential links with disease outcomes.

## Discussion

In this study, we show that CD4^+^ and CD8^+^ T-cell responses in DENV infection were highly enriched in the skin where these cells displayed phenotypic and functional features of T_RM_ cells. Responding CD4^+^ and CD8^+^ T-cells in the skin displayed increased expression of skin residency markers CD69, CD103 and of PD-1 and granzyme B, compared to their blood counterparts. ScRNA-seq of activated CD8^+^ T-cells in the skin and blood showed distinct gene expression profiles of these cells. Skin CD8^+^ T-cells expressed transcriptional signatures of human T_RM_ cells(*18*), supporting their differentiation into T_RM_ cells during acute infection. The magnitude of CD8^+^ T-cell responses in the skin mirrored that of their blood counterparts, while skin CD4^+^ T-cell frequencies were independent from those of matched blood CD4^+^ T-cells, suggesting a differential regulation of CD4^+^ and CD8^+^ T-cell responses between these compartments. Outpatients, who did not require hospitalization, displayed a higher magnitude of skin and blood CD8^+^ T-cell responses and increased expression of skin T-cell related cytokines compared to inpatients. This study provides novel insights into the biology and dynamics of T_RM_ cells in a human acute viral infection and suggests a protective role of both skin and blood CD8^+^ T-cells in dengue.

We previously showed in dengue patients that circulating DENV-specific T-cells express high levels of CLA while human cytomegalovirus (CMV)-specific T-cells that are “bystander activated” during DENV infection lack expression of CLA(*7, 24*). These studies showed that co-expression of Ki-67 and CLA distinguishes circulating DENV-specific T-cells from bystander activated T-cells. From these studies it remained unclear whether Ki-67^+^ CLA^+^ DENV-specific T-cells differentiate into T_RM_ cells in the skin and whether T-cells responding in the skin and blood are phenotypically and functionally distinct.

Here we show that CD4^+^ and CD8^+^ Ki-67^+^ CLA^+^ T-cells are highly enriched in the skin compared to the blood compartment, suggesting their preferential localization within this site. At days 7-10 from illness onset (corresponding approximately to days 11-14 from infection), responding CD8^+^ T-cells in the skin displayed transcriptional and phenotypic features of T_RM_ cells. These cells expressed a core T_RM_ gene signature which includes: upregulation of CD69 and downregulation of S1PR1 and KLF2 which collectively lead to T-cell retention within tissues; upregulation of adhesion molecules (CD103, CD49a) and PD-1; modulation of chemokine receptor expression (CXCR6, CX3CR1) and downregulation of transcriptional factors Eomes and T-bet(*18*). Skin CD8^+^ T-cells displayed increased expression of genes encoding for granzyme B, perforin and granulysin. Consistently, flow cytometry analyses showed strikingly higher frequencies of granzyme B^+^ CD4^+^ and CD8^+^ T-cells in the skin compared to those in the blood of the same patients, supporting a strong cytotoxic capacity of skin T-cells in DENV infection. These data are consistent with previous studies in healthy individuals showing that despite the low expression of perforin and granzymes by skin CD8^+^ T_RM_ cells in steady state, these cells can efficiently induce expression of these molecules upon cytokine or TCR stimulation(*26*).

Previous studies in mice showed that virus-specific CD8^+^ T-cells are highly polyclonal with >1000 different clonotypes responding to a single immunodominant virus epitope(*27*). This polyclonal response, which comprises of T-cells with differing levels of TCR avidity, ensures protection towards potential pathogen escape mutants(*28*). Similarly in humans, CD8^+^ T-cells targeting the immunodominant CMV NLV epitope display high clonality, with >5,000 different clonotypes targeting the same epitope(*29*). Here we show that the clonotype diversity of activated CD8^+^ T-cells in DENV infection was low/moderate and similar in skin and blood compartments, with approximately 150-350 unique clonotypes identified per patient sample. Patient 2 displayed the lowest clonotype diversity and was experiencing a secondary DENV infection, while patients 1 and 3 were primary dengue cases. While a lower clonotype diversity in secondary DENV infection is consistent with a preferential reactivation of memory T-cells generated in the primary infection, further studies on a higher number of patients are needed to study clonotypic diversity in primary and secondary DENV infection and address potential links with disease outcomes. In patients 2 and 3 we detected some clonotype sharing between skin and blood (approximately 7%), suggesting recirculation of responding CD8^+^ T-cells between these two compartments. However, the limited proportion of clonotype sharing between skin and blood suggests some degree of compartmentalization of the CD8^+^ T-cell response and independent priming of these cells at different sites. These findings are consistent with the nature of DENV infection which initiates in the skin but then spreads systemically potentially affecting multiple organs(*30*).

A limitation of our analyses is the small number of patient samples included in the scRNAseq and the peptide-HLA pentamer flow cytometry analyses. The latter was due to difficulties in identifying dengue patients expressing HLA-A*11:01 (approximately 30% of patients in this cohort) and HLA-A*11:01^+^ patients that displayed a detectable response to the two epitopes analysed (NS3_1608-1617_ and NS5_2610-2618_), despite these being the most immunodominant epitopes restricted to this HLA type(*24, 25*). This study was not powered to compare T-cell responses in patients infected with different DENV serotypes. The predominant serotype of infection in this cohort was DENV 2 (47%) followed by DENV 3 (25%) with DENV 1 and DENV 4 infections being less common (4%). For 25% of patients, we were unable to determine the serotype of infection as their RNAemia was below the detection limit at enrolment.

While the majority of the T-cell response in dengue comprised of Ki-67^+^CLA^+^ cells, we detected three distinct clusters of Ki-67^+^ CLA^−^ CD8^+^ T-cells, two of which were highly enriched in the blood compared to the skin compartment, while a small cluster of cells (approximately 3% of CD8^+^ T-cells) was equally represented in the two compartments. Ki-67^+^ CLA^−^ CD8^+^ T-cells may represent bystander activated CD8^+^ T-cells that were primed in sites different from the skin or migrated from the skin to the blood and lost CLA expression. Previous studies showed that virus infection may trigger activation of T-cells specific for persistent herpesviruses (HCMV and Epstein-Barr virus) and this is likely mediated by IL-15 produced during the infection(*7, 31*). In this study, due to limiting number of cells present in the skin blisters, we were unable to address the antigen-specificity of these cells and the signals driving their activation.

Our study shows that Ki-67^+^ CLA^+^ CD8^+^ T-cells which phenotypically and functionally mirror DENV-specific CD8^+^ T-cells identified using peptide-HLA tetramers(*24, 25, 32*), were present at higher frequencies in the skin and blood of patients with milder disease, who did not require hospitalization. Consistently, cytokines produced by effector T-cells (IL-2, IL-17) or cytokines/chemokines that drive differentiation and recruitment of these cells were increased in skin blisters from outpatients compared to inpatients. IL-17 is expressed by human CD4^+^ T_RM_ and CD8^+^ T_RM_ cells and was also shown to be induced in T-cells of healthy volunteers by mosquito saliva in a human challenge study(*33*).

In summary, this study shows that a large proportion of the T-cell response to DENV resides within the skin and suggests a protective role of T_RM_ and circulating CD8^+^ T-cells. Our findings warrant evaluation of vaccine strategies that elicit priming of CD8^+^ T_RM_ cells.

## Materials and methods

### Study design and study participants

Dengue patients (N=73) and healthy volunteers (N=10) included in this study were recruited at Tan Tock Seng Hospital and National Centre for Infectious Diseases between June 2019 and April 2023, Singapore, after written informed consent following Institutional Review Board (IRB) approval [National Healthcare Group (NHG) Domain Specific Review Board (DSRB) Reference: 2018/00874]. Clinical and demographic details of patients and healthy volunteers are summarised in **Table I**. Dengue disease classification was performed by trained clinicians at the Tan Tock Seng Hospital/National Centre for Infectious Diseases and was based on the WHO 2009 definition(*34*).

Inclusion criteria for dengue patients include: a confirmed diagnosis of dengue fever (NS1-positive and/or serology-positive), no other diagnosis supporting the febrile episode and ≥21 years of age. Inclusion criteria for healthy volunteers include: not being diagnosed with dengue and with no serological evidence of past DENV infection, and ≥21 years of age. Exclusion criteria for dengue patients and healthy volunteers include: pregnancy and breast-feeding (for women), having asthma and existing skin conditions or being immunocompromised. Whole blood was collected at visits 1 and 2 (days 3-5 and 7-10 from fever onset) in K3EDTA tubes and skin suction blisters were raised at visit 2 in the same volunteers.

### DENV IgG

The Panbio Dengue IgG Indirect ELISA test was performed using plasma samples from dengue patients and healthy individuals according to the manufacturer’s protocol (Abbott, Cat No: 01PE30). The absorbance was read using the Varioskan LUX multimode microplate reader. Primary and secondary DENV infection were defined based on serum DENV IgG levels at days 3-5, as defined previously(27).

### Viral RNA extraction & DENV serotype analyses

The extraction of viral RNA from plasma was carried using the QIAamp Viral RNA Mini Kit (Qiagen), following the manufacturer’s protocol. DENV serotype of infection was determined using the Center for Disease Control and Prevention (CDC) DENV-1–4 real-time RT-PCR (qRT-PCR) Assay protocol with the Invitrogen SuperScript® III One-Step Quantitative RT-PCR kit (Invitrogen, Cat No: 11732-088) on the Roche LightCycler 96. The primers and probes used are listed in **Table S2**.

### DNA extraction & HLA-A typing

The DNA extraction protocol was adapted and modified from the Qiagen AllPrep DNA/RNA Mini Kit. Briefly, 0.5mL of clotted blood was mixed with RLT buffer and β-mercaptoethanol. The solution was vortexed, then incubated at room temperature for 5 minutes. 600μL of the blood-RLT buffer solution was transferred to an AllPrep DNA spin column, and spun down at 10, 000rpm for 1 minute at 18°C. The flowthrough was discarded, and the process repeated until all the blood-RLT buffer solution had been spun down. The spin column was then washed twice with 500μL Buffer AW1 at 10,000rpm for 1 minute at 18°C, discarding the flowthrough for each wash. The spin column was washed once with 500μL Buffer AW2 and spun down at 14,000rpm for 2 minutes at 18°C, discarding the flowthrough. The spin column was spun down once more at 14,000rpm to dry the column. 50μL of Buffer EB, pre-heated to 70°C, was added to the spin column and let sit for 2 minutes, before it was centrifuged at 10,000rpm for 1 minute. The process was repeated once more without discarding the flowthrough until the final volume of DNA eluted was approximately 100μL. The DNA concentration was measured using a NanoDrop 2000. The HLA-A typing assay was adapted and modified from a previous study(*35*). For every reaction, a mix of 0.5μL of dNTP mix (Promega, Cat No: U1511), 2.5μL of (NH_4_)SO_4_-Cl_2_ buffer, 2μL of MgCl_2_, 0.25μL of *Taq* DNA polymerase (Thermo Scientific, Cat No: EP0402) and 2μL of each primer (A1101-AL#6 sequence: CGG AAT GTG AAG GCC CAG & A1101-AL#1 sequence: TCT CTG CTG CTC CGC CG) was prepared to a total volume of 9.25μL. 100ng of the DNA sample of interest or positive control was added to the mix, and nuclease-free water was topped up to a total volume of 25μL per PCR reaction. As a negative control, nuclease-free water was added in place of DNA. Thermocycling parameters were as follows: Stage 1, 96°C for 15 minutes for 1 cycle; Stage 2, 5 cycles of 96°C for 1 minute, 66°C for 1 minute and 72°C for 1 minute; Stage 3, 40 cycles of 96°C for 1 minute, 56°C for 1 minute, 72°C for 1 minute, and final hold at 4°C. PCR products were analysed by agarose gel electrophoresis.

### Blood processing and PBMC isolation

Whole blood was collected in K3EDTA tubes. 1mL of the whole blood was aspirated into a 1.5ml Eppendorf tube. The tube is spun down in a benchtop centrifuge at 3000rpm for 5 minutes. The clotted blood was used for DNA extraction for HLA-typing. PBMCs were isolated from the remaining whole blood by Ficoll-Paque gradient centrifugation as previously described(*36*). Freshly isolated PBMCs were used for flow cytometry and RNA sequencing assays on the day of collection.

### Skin blister induction

The skin blister suction method was adapted from previous studies(*7, 21*). Using a clinical pump connected to a chamber, blisters were induced on the forearm of the study volunteers. A negative pressure of 25-40kPa (200-300mmHg) below atmospheric pressure was applied for 2-4 hours until a blister was formed. After 18-24 hours, the blister fluid was aspirated using a sterile 23G needle and 2mL syringe, and transferred to a 1.5mL Eppendorf tube. Skin cells were recovered by centrifugation at 3000 rpm for 10 minutes at 4°C. The blister fluid supernatant was cryopreserved and used for Olink Target 48 assay. The skin cell pellet was re-suspended in 200μL of AIM-V (Gibco, Cat No: 12055091) + 2% Human Serum medium for analysis. Freshly isolated cells from the skin blister were used for flow cytometry and RNA sequencing assays on the day of collection.

### Flow cytometry

Staining was performed on freshly isolated matched PBMCs and skin cells on the day of collection. Cells were stained with LIVE/DEAD™ Fixable Blue Stain for 10 minutes in the dark at room temperature. Cells were then stained with antibodies targeting surface markers in staining buffer (PBS + 1% bovine serum albumin, BSA), and with phycoerythrin (PE)-labelled DENV pentamers, for 20 minutes on ice. For panels with Ki-67 & granzyme B staining, cells were suspended with 100μL of eBioscience FoxP3 Fixation/Permeabilization solution (Invitrogen, Cat No: 00-5523-00) for 45 minutes on ice, before washing with Permeabilization buffer (Invitrogen, Cat No: 00-5523-00). Cells were then stained with antibodies targeting intracellular markers for 30 minutes on ice, before acquisition on the BD LSRFortessa™. The list of antibodies and peptide-HLA pentamers used is provided respectively in **Tables S3** and **S4**. Flow cytometry data was analysed on FlowJo Version 10; plugins include DownSample, UMAP, PhenoGraph and ClusterExplorer.

### Olink Target 48 analysis

Analyses of the following 45 cytokines/chemokines was performed in skin blister fluid supernatant according to the manufacturer’s instructions using the Olink Target 48 Cytokine panel: IL-18, HGF, CCL19, CCL2, MMP12, LTA, FLT3LG, TNF, IL-17A, IL-2, IL-17F, CSF3, IL-1β, OLR1, TNFSF12, CXCL10, VEGFA, IL33, TSLP, IFN-γ, CCL4, TGFA, IL-13, CXCL8, CCL8, IL-6, CCL13, CSF2, CCL7, IL-4, TNFSF10, OSM, MMP1, EGF, IL-7, IL-15, CSF1, CXCL9, CXCL11, IL-17C, CXCL12, CCL11, IL-10, CCL3, EBI3, IL-27. 45 oligonucleotide conjugated antibody pairs were hybridized with skin blister fluid and incubated overnight at 4°C. Oligonucleotides brought into proximity hybridize, with extension facilitated by a DNA polymerase that results in the formation of a target-specific double stranded DNA barcode. The 45 unique DNA barcode sequences were then amplified by PCR before being quantified using the Olink Signature Q100.

### Single-cell RNA sequencing

CD8^+^ CD38^+^ HLA-DR^+^ cells from matched PBMCs and skin blister cells of dengue patients were sorted into AIM-V + 2% Human Serum on the BD FACSAria™ III Sorter following staining with antibodies targeting surface markers. Single-cell RNA sequencing of the sorted cells was carried out according to the 10X Genomics’ Chromium Next GEM Single Cell 5’ Reagent Kits v2 protocol. Briefly, sorted cells were mixed with 10x barcoded gel beads and loaded onto Chromium Next GEM Chip K, after which the assembled chip was run on the Chromium Controller X to generate GEMs (1000263 Chromium Next GEM Single Cell 5’ Kit v2, 1000286 Chromium Next GEM Chip K Single Cell Kit & 1000190 Library Construction Kit, 10X Genomics). Post GEM-RT cleanup was then carried out, followed by cDNA amplification according to manufacturer’s protocol. Subsequently, V(D)J amplification from cDNA, V(D)J library construction as well as 5’ gene expression library construction were carried out according to manufacturer’s protocol (1000252 Chromium Single Cell Human TCR Amplification Kit, 10X Genomics). Quality control was performed on the Agilent 2100 Bioanalyzer G2938C. Libraries were sequenced paired-end 150 base pairs on the Illumina HiSeq X System (20,000 read pairs per cell).

### Single cell RNA-seq data processing and analysis

Raw sequencing data were processed using the Cell Ranger multi pipeline (v7.1.0, 10x Genomics). The filtered gene-barcode matrix of unique molecular identifiers (UMI) counts was then analyzed with Seurat V3(*37*) for quality control, normalization, dimensional reduction, integration, clustering, and visualization. Quality control criteria were: (1) total UMI count between 3,000 and 30,000, (2) minimal number of detected genes > 1000, (3) mitochondrial gene percentage <15% and (4) number of cells expressing a gene > 8. The count matrix was log-normalized, and the top 2,000 most variable genes were identified for dimensional reduction.

The count matrix was scaled and the top 20 dimensions from the principal component analysis (PCA) were used for the uniform manifold approximation and projection (UMAP). The top PCs were chosen using the Elbow plot method. Cell clusters were identified by the shared nearest neighbor (SNN) method using the Louvain algorithm with the resolution of 1.2 and the top 20 PCA dimensions. Cell clusters of CD8^+^ T-cells were annotated using known markers of T-cells as well as referencing cell clustered annotated using SingleR, which compared the transcriptome of each cell cluster to various reference datasets (Human primary cell atlas, Database Immune Cell Expression Data and Monaco immune data). For the final analysis, cell clusters were merged into six groups (Naïve, Cell cycle, Blood, Skin, Mait-like and Memory-like).

### scTCR-seq data processing and analysis

Sequencing data were processed using the Cell Ranger VDJ pipeline (v7.1.0, 10x Genomics) with default settings. Briefly, Cell Ranger aligned TCR reads to the GRCh38 reference genome, determined consensus TCR sequences and identified contigs and the corresponding CDR3 regions and V, D, J, C genes. TCR clonotypes represented by CDR3 amino acid sequences and V gene usages were analysed using scRepertoire(*38*).

### Statistical analysis

GraphPad Prism 10 (GraphPad Software Inc, California, USA) was used for statistical analyses of flow cytometry data. Wilcoxon matched-pairs sign rank test was used for comparisons between blood and skin. Mann-Whitney t-test was used for comparisons between two groups (dengue vs healthy, and primary vs secondary patients, outpatients vs inpatients) and One-way ANOVA with Benjamini-Hochberg FDR correction for multiple comparisons was used for comparison of more than two groups. Statistical analyses for the Olink data was performed by One-way ANOVA using Partek Genomic Suite Analysis Version 7 software. For all analyses p-values are defined as follows: *p ≤0.05, **p ≤0.01, ***p ≤0.005, ****p ≤0.0001.

## Acknowledgments

We are grateful to all patients and healthy volunteers who took part in this study. Thank you to Tan Bee Har, Junni Tang and all the medical and nursing staff who helped with participant recruitment. This work was supported by an Open Fund–Individual Research Grant, National Medical Research Council, Singapore to LR (NMRC/OFIRG/0049/2017).

MG is supported by the Academy of Medical Sciences and the Springboard Award scheme funders: the Wellcome Trust, the Government Department of Business, Energy and Industrial Strategy and the British Heart Foundation and Diabetes UK (SBF007\100173 to LR).

## Author contributions

Conceptualization: LR.

Methodology: LR, NZH, CKW, TLT.

Investigation: NZH, CKW, EOZ, VC, TBH, JT.

Visualization: NZH, JSGO, MG.

Funding acquisition: LR, OEE.

Project administration: LR, OEE, DLCB.

Supervision: LR, OEE.

Writing – original draft: LR and NZH.

Writing – review & editing: all authors.

## Data and materials availability

Flow cytometry and scRNAseq data will be deposited in the relevant repositories upon acceptance of the paper. This paper does not report original codes. Any additional information required to re-analyze the data reported in this paper is available from the lead contact upon request.

