## Supplementary material for "Dengue infection elicits skin tissue-resident and circulating CD8^+^ T-cells associated with protection from hospitalization"

### Supplementary Materials

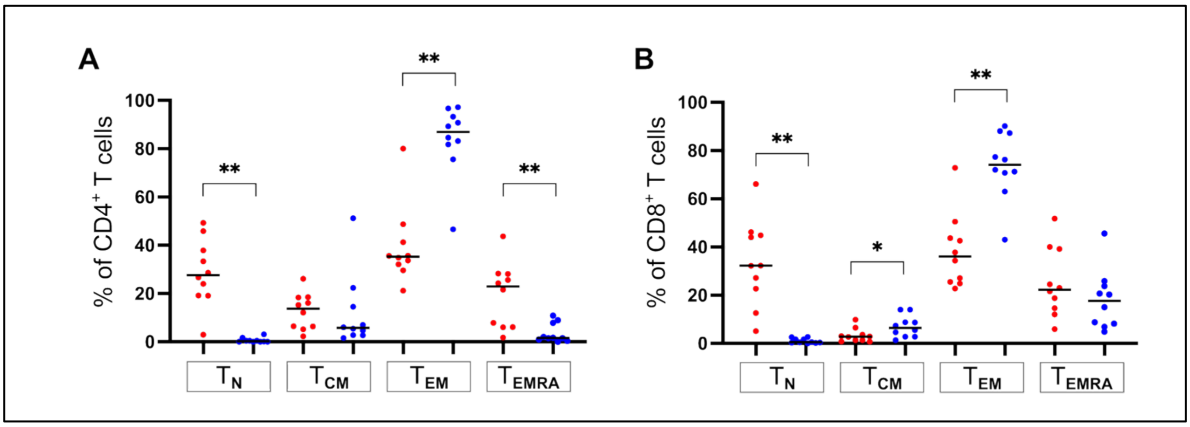


**Figure S1. Frequencies of blood and skin T-cell subsets in healthy volunteers.** T-cell subsets defined by CCR7 and CD45RA expression: $T_N$ (naïve): CCR7+CD45RA+; $T_{CM}$ (T central memory): CCR7+CD45RA-; $T_{EM}$ (T effector memory): CCR7-CD45RA-; $T_{EMRA}$ (T effector memory re-expressing CD45RA): CCR7-CD45RA+. Datapoints for blood and skin samples for each participant are shown respectively in red and blue. Statistics were determined by Wilcoxon matched-pairs sign rank test between blood and skin.

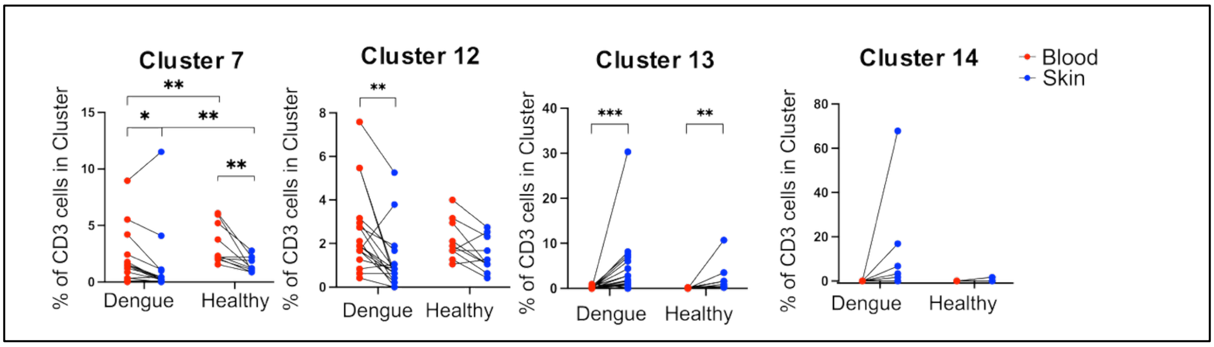


**Figure S2. Skin and blood T-cells are largely distinct.** Frequencies of cells within clusters defined by phenograph in Fig. 3 in the skin and blood of dengue patients and healthy volunteers. Statistics were calculated by Wilcoxon matched-pairs sign rank test between blood and skin, and Mann-Whitney t-test between healthy volunteers and patients.

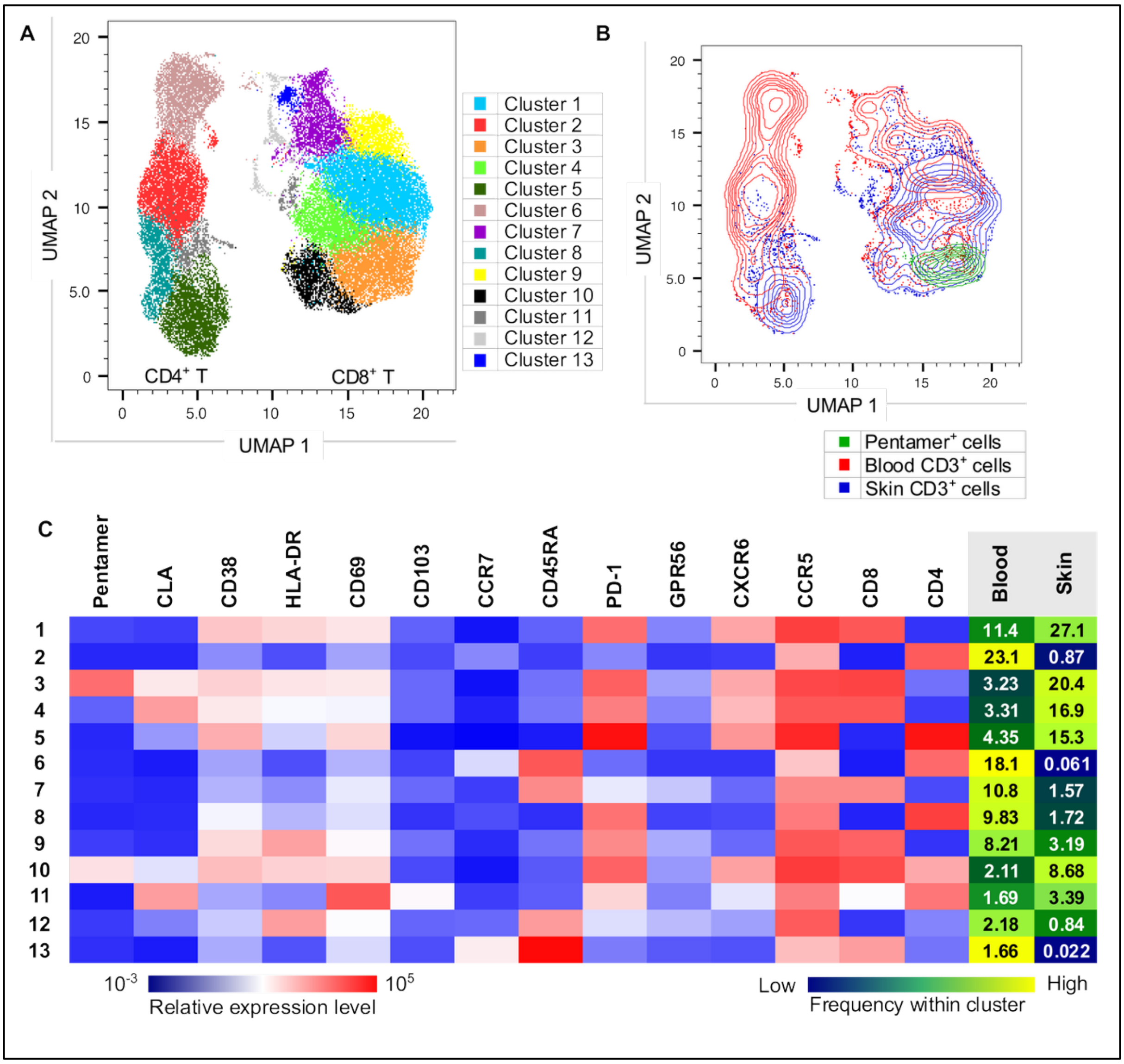


**Figure S3. DENV pentamer+ CD8+ T-cells are enriched in the skin. (A)** UMAP plot with phenograph clustering is shown for skin and blood T-cells of a dengue patient. **(B)** Skin (blue) and blood (red) CD3+ T cells, and manually-gated pentamer+ DENV-specific CD8+ T cells (green) cells are shown. **(C)** Mean fluorescence intensity (MFI) of the analysed markers for each cluster are displayed is shown in a heatmap. Frequencies of each phenograph cluster within the blood and skin are displayed in the respective columns.

**Table S1:** Details for the patients included in the ScRNA-seq analyses. Indicated are the number of activated CD8$^{+}$ T-cells sorted from each patient sample.

| Patient | Days from fever onset | Age | Infection | Rash | Hospitalization | Bleeding | Blood- No. of CD38$^{+}$HLA-DR$^{+}$ CD8+ T cells | Skin- No. of CD38+HLA-DR+ CD8+ T cells |
|---|---|---|---|---|---|---|---|---|
| P1 | 10 | 29 | Primary | Yes | Outpatient | No | 1700 | 803 |
| P2 | 7 | 26 | Secondary | Yes | Outpatient/ Inpatient | Yes | 1700 | 612 |
| P3 | 8 | 31 | Primary | No | Inpatient | Yes | 2000 | 1510 |

**Table S2:** Primers and probes used for the Center for Disease Control and Prevention (CDC) DENV-1–4 RT-PCR Assay.

| | Primers | | Probes | |
|---|---|---|---|---|
| DENV 1 | F | 5’ CAA AAG GAA GTC GTG CAA TA 3’ | D1-FAM-Probe | 5’ /6-FAM/ CAT GTG GTT /ZEN/ GGG AGC ACG C /3IABkFQ/ 3’ |
| | C | 5’ CTG AGT GAA TTC TCT CTA CTG AAC C 3’ | | |
| DENV 2 | F | 5’ CAG GTT ATG GCA CTG TCA CGA T 3’ | D2-HEX-Probe | 5’ /HEX/ CTC TCC GAG /ZEN/ AAC AGG CCT CGA CTT CAA /3IABkFQ/ 3’ |
| | C | 5’ CCA TCT GCA GCA ACA CCA TCT C 3’ | | |
| DENV 3 | F | 5’ GGA CTG GAC ACA CGC ACT CA 3’ | D3-TexRd-Probe | 5’ /TexRd-XN/ ACC TGG ATG TCG GCT GAA GGA GCT TG /3IAbRQSp/ 3’ |
| | C | 5’ CAT GTC TCT ACC TTC TCG ACT TGT CT 3’ | | |

| DENV 4 | F | 5’ TTG TCC TAA TGA TGC TGG TCG 3’ | D4 Cy5-Probe | 5’ /Cy5/ TTC CTA CTC /TAO/ CTA CGC ATC GCA TTC CG /3IAbRQSp/ 3’ |
|---|---|---|---|---|
| | C | 5’ TCC ACC TGA GAC TCC TTC CA 3’ | | |

**Table S3:** Flow cytometry antibodies used.

| | Antibody | Clone | Brand & Catalogue number | Staining volume (µL in 50µL) |
|---|---|---|---|---|
| 1 | BUV737 Mouse Anti-Human CD279 (PD-1) | EH12.1 | BD Horizon, 612791 | 3 |
| 2 | BUV395 Mouse Anti-Human CD45 | HI30 | BD Horizon, 563792 | 3 |
| 3 | V500 Mouse Anti-Human CD3 | UCHT1 | BD Horizon, 561416 | 3 |
| 4 | BV711 Mouse Anti-Human CD38 | HIT2 | BD Horizon, 563965 | 2.5 |
| 5 | BV650 Mouse Anti-Human CD314 (NKG2D) | 1D11 | BD Horizon, 563408 | 5 |
| 6 | BUV737 Mouse Anti-Human CD195 (CCR5) | 2D7/CCR5 | BD Horizon, 565293 | 5 |

| | | | | |
|---|---|---|---|---|
| 7 | APC-Cy7 Mouse Anti-Human CD8 | SK1 | BD Pharmingen, 557834 | 3 |
| 8 | Alexa Fluor® 700 Mouse anti-Human CD197 (CCR7) | 150503 | BD Pharmingen, 561143 | 3 |
| 9 | PE Mouse Anti-Human CD45RO | UCHL1 | BD Pharmingen, 555493 | 3 |
| 10 | PE-Cy7 Mouse Anti-Human HLA-DR | L243 | BD, 335795 | 2.5 |
| 11 | Brilliant Violet 785 anti-human CD69 | FN50 | Biolegend, 310932 | 5 |
| 12 | Brilliant Violet 650™ anti-human HLA-DR | L243 | Biolegend, 307650 | 1.3 |
| 13 | BV605 Mouse Anti-Human CD45RA | HI100 | Biolegend, 562886 | 0.5 |
| 14 | FITC anti-human/mouse Cutaneous Lymphocyte Antigen (CLA) | HECA-452 | Biolegend, 321306 | 10 |
| 15 | PE/Cyanine7 anti-human GPR56 | CG4 | Biolegend, 358206 | 2.5 |
| 16 | PE/Dazzle™ 594 anti-human CD4 Antibody | A161A1 | Biolegend, 357412 | 1 |

| | | | | |
|---|---|---|---|---|
| 17 | Brilliant Violet 421™ anti-human CD279 (PD-1) Antibody | EH12.2H7 | Biolegend, 329920 | 5 |
| 18 | Alexa Fluor® 647 anti-human CD186 (CXCR6) Antibody | K041E5 | Biolegend, 356008 | 2 |
| 19 | Brilliant Violet 421™ anti-human Ki-67 Antibody | Ki-67 | Biolegend, 350506 | 5 |
| 20 | CD103 (Integrin alpha E) Monoclonal Antibody PerCP-eFluor710 | Ber-ACT8 | eBioscience, 46-1037-42 | 2.5 |
| 21 | APC anti-human Granzyme B Antibody | GB11 | Invitrogen, GRB05 | 5 |
| 22 | LIVE/DEAD™ Fixable Blue Dead Cell Stain Kit, for UV excitation | | Life Technologies, L23105 | 1:1000 |

**Table S4:** List of PE-conjugated peptide-HLA pentamers purchased from ProImmune Limited and the sequence for each peptide are shown.

| | Sequence | Epitope | Staining volume (µL in 50µL) |
|---|---|---|---|
| **DENV 1, HLA-A*11:01** | GTSGSPIVNR | NS3 1608-1617 | 10µL each |
| | ATYGWNLVK | NS5 2610-2618 | |
| **DENV 2, HLA-A*11:01** | GTSGSPIIDK | NS3 1608-1617 | |
| | STYGWNLVR | NS5 2610-2618 | |
| **DENV 3, HLA-A*11:01** | GTSGSPIINR | NS3 1608-1617 | |
| | STYGWNIVK | NS5 2610-2618 | |
| **DENV 4, HLA-A*11:01** | GTSGSPIINR | NS3 1608-1617 | |
